# Predicting mutation-disease associations through protein interactions via deep learning

**DOI:** 10.1101/2024.08.06.606730

**Authors:** Xue Li, Ben Cao, Jianmin Wang, Xiangyu Meng, Shuang Wang, Yu Huang, Enrico Petretto, Tao Song

**Author notes:** **Corresponding Author** Tao Song – School of Computer Science and Technology, China University of Petroleum (East China), Qingdao, Shandong, China;, Enrico Petretto - Centre for Computational Biology, Duke-NUS Medical School, Singapore. Xue Li and Ben Cao are joint first authors.

## Abstract

Disease is one of the primary factors affecting life activities, with complex etiologies often influenced by gene expression and mutation. Currently, wet-lab experiments have analyzed the mechanisms of mutations, but these are usually limited by the costs of wet experiments and constraints in sample types and scales. Therefore, this paper constructs a real-world mutation-induced disease dataset and proposes Capsule networks and Graph topology networks with multi-head attention (CGM) to predict the mutation-disease associations. CGM can accurately predict protein mutation-disease associations, and in order to further elucidate the pathogenicity of protein mutations, we also verified that protein mutations lead to protein structural alterations by Swiss-model, which suggests that mutation-induced conformational changes may be an important pathogenic factor. Limited by the size of the mutated protein dataset, we also performed experiments on benchmark and imbalanced datasets, where CGM mined 22 unknown protein interaction pairs from the benchmark dataset, better illustrating the potential of CGM in predicting mutation-disease associations. In summary, this paper curates a real dataset and proposes CGM to predict the protein mutations-disease associations, providing a novel tool for further understanding of biomolecular pathways and disease mechanisms.

## 1 INTRODUCTION

DNA sequence mutations can lead to abnormal gene expression, and can also affect protein function. Some mutations can directly alter protein solubility ^1^, thereby changing protein function and structure, leading to disease onset ^2, 3^. For example, mutations in the PSEN1 and APOE ε4 genes lead to abnormalities in presenilin-1 function and APOE structure ^4, 5^, triggering Alzheimer’s disease. Mutations in the TP53 and BRCA1/BRCA2 lead to changes in the tumor suppressor protein p53 ^6^ and DNA repair proteins ^7^, increasing cancer risk. Additionally, proteins may also undergo post-translational changes due to a number of factors. Mutations in amyloid precursor proteins lead to abnormal deposition of amyloid beta (Aβ), a major pathological feature in familial Alzheimer’s disease cases ^8^. Collagen protein mutations may promote tumor development by disrupting protein folding, leading to functional and structural changes or loss ^9^. Furthermore, mutations in proteins such as SHP2 phosphatase, associated with Noonan syndrome with multiple lentigines, result in reduced enzymatic activity and structural differences ^1^. Therefore, protein-coding mutations can be significant contributors to the onset and progression of diseases. This is because mutations can alter the intrinsic properties of proteins ^10^, inducing new protein interactions or weakening existing ones, thus reshaping protein-protein interaction (PPIs) networks ^11^. Changes in PPI networks can affect mechanisms such as biological signal transduction, ultimately leading to disease.

To elucidate how mutated proteins alter PPIs networks leading to disease mechanisms, various experimental assays in model organisms and cellular systems are commonly used. For example, Wang et al. through analysis of serial clinical blood samples, proposed that the downregulation effect of the DACT1^M576K^ mutation on the DVL2 protein may potentially lead to congenital heart disease ^12^. In 2022, Mo et al. employed methods like Bidirectional Encoder Representations from Transformers (BERT) to construct a reconstructed protein network involving four cancer-suppressing genes including BRAF^V600E^ and others. This study demonstrated the relevance new protein interactions to disease, where mutations in cancer-suppressing proteins not only lose existing interactions, but also induce new protein interactions, thereby promoting tumor initiation and progression. However, these studies often focus on specific protein mutation sites, reducing the weight of mutation on global protein sequences that determine protein structure ^13^. It is noteworthy that protein mutations can cause structural and functional changes ^14^, which are common triggers for diseases ^15^. Experimental validation of mutation-disease associations is typically feasible only in model animals or cellular tissues and require significant funding and time. Moreover, most experiments are limited as they are designed for specific cell types and proteins, lacking universality and scalability. When studying new protein-coding mutations, existing experimental processes and methods often face challenges due to optimization of experimental protocols, which can be extensive and are difficult generalize to multiple diseases. In silico methods, however, offer high scalability and adaptability, making them a feasible alternative for constructing predictive models to explore mutation-disease associations.

Deep learning is currently a useful method for constructing predictive models, particularly excelling in genomics and proteomics^16-18^. A deep learning model used to substantially improve gene expression accuracy, named Enformer^18^, was able to predict enhancer-promoter interactions directly from DNA sequences. Wang et al. used a deep learning model to explore the conformational ensembles of protein−protein complexes through MD simulations. Current protein prediction models are constrained by data volume ^19^ and structural prediction accuracy ^20^, and primarily rely on sequence features. Therefore, in PPIs prediction models, embedding sequence features is a classical representation scheme ^19, 21-23^, such as Auto Covariance (AC), Local Descriptors (LD), Multi-scale Continuous and Discontinuous (MCD), Conjoint Triad (CT), Pseudo-Amino Acid Composition (PseAAC). However, these descriptor-based representation methods, while diverse, may not be comprehensive enough, especially for mutations with small impact within a single protein sequence. Thus, protein pre-training models supported by computational resources are needed ^24-26^, which can learn protein features and patterns from large amounts of protein sequence data. Although pre-training models based on large-scale data knowledge can help overcome feature representation bottlenecks, they often overlook overall protein network structural information, hence limiting our ability to assess potential unknown protein-protein interactions. Zhu et al. ^27^, building upon descriptors, introduced the SGAD model based on Structural Deep Network Embedding to PPIs, augmenting multiple protein sequence descriptors, as well as topological features and information flow of the PPI network,, identifying potential interaction pairs like FZD5-WDR1 and RHOA-NR2F6 in cancer. Subsequently, Lei et al. ^28^ proposed a deep learning framework for multi-level peptide-protein interaction prediction (CAMP) including binary peptide-protein interaction prediction and corresponding peptide binding residue identification. These methods have discovered novel protein interaction pairs, potentially predicting abnormal protein interactions relevant to disease etiology or progression. However, constrained by availability of experimental data ^29^, these methods cannot infer the mutation-disease associations of mutated proteins within a wider protein network. Thus, constructing reliable and extensive experimental datasets for mutated proteins and interaction prediction models is necessary.

In this paper, we construct a mutation-induced disease dataset focused on mutations associated with cancer and cardiac disease, and propose a Capsule and Graph topology networks with Multi-head attention (CGM) model to predict novel protein interaction induced by mutations, and assess the impact of novel PPIs on disease. The results demonstrate that CGM achieves high-quality predictions of mutated protein interactions in real datasets. To further explore the pathogenic mechanisms of mutated proteins, we also verified that the predicted protein mutations lead to structural changes by Swiss-model. Additionally, validation on benchmark datasets and imbalanced datasets related to diabetes, cardiac disease, Alzheimer’s disease (AD), Parkinson’s disease (PD), and cancer further support the wide applicability and robustness of CGM. We also report 22 previously unknown protein interaction pairs, which further illustrates CGM’s capability to learn features of mutated protein information. In summary, by constructing a dataset of proteins mutated and proposing a deep learning-based prediction model, this paper strengthen the paradigm that structural alterations due to protein mutations is likely to be one of the main causes of disease, providing a new perspective for understanding PPI network changes in disease.

## 2 MATERIALS AND METHODS

To address the limitations of current mutation-disease association predictions due to data and wet-lab experiment scale constraints, this paper first constructs a mutation-induced disease dataset of mutated proteins (Figure 1A) and then an interaction prediction model (Figure 1C). With the real dataset and the prediction model, this paper elucidates the mechanisms by which mutations cause PPI network reorganization can lead to disease (Figure 1B). Initially, relevant literature was collected through keyword screening and retrieval, ensuring data validity and reproducibility. Subsequently, the dataset was curated to include only proteins with mutations and their interacting proteins (without mutation), and mutated proteins are annotated for both wild type and mutant types. Finally, the CGM model is designed to predict on mutation-based protein interactions, thereby assessing the impact of novel PPIs caused by mutations on diseases, to yield mutation-disease association predictions.

**Figure 1.**
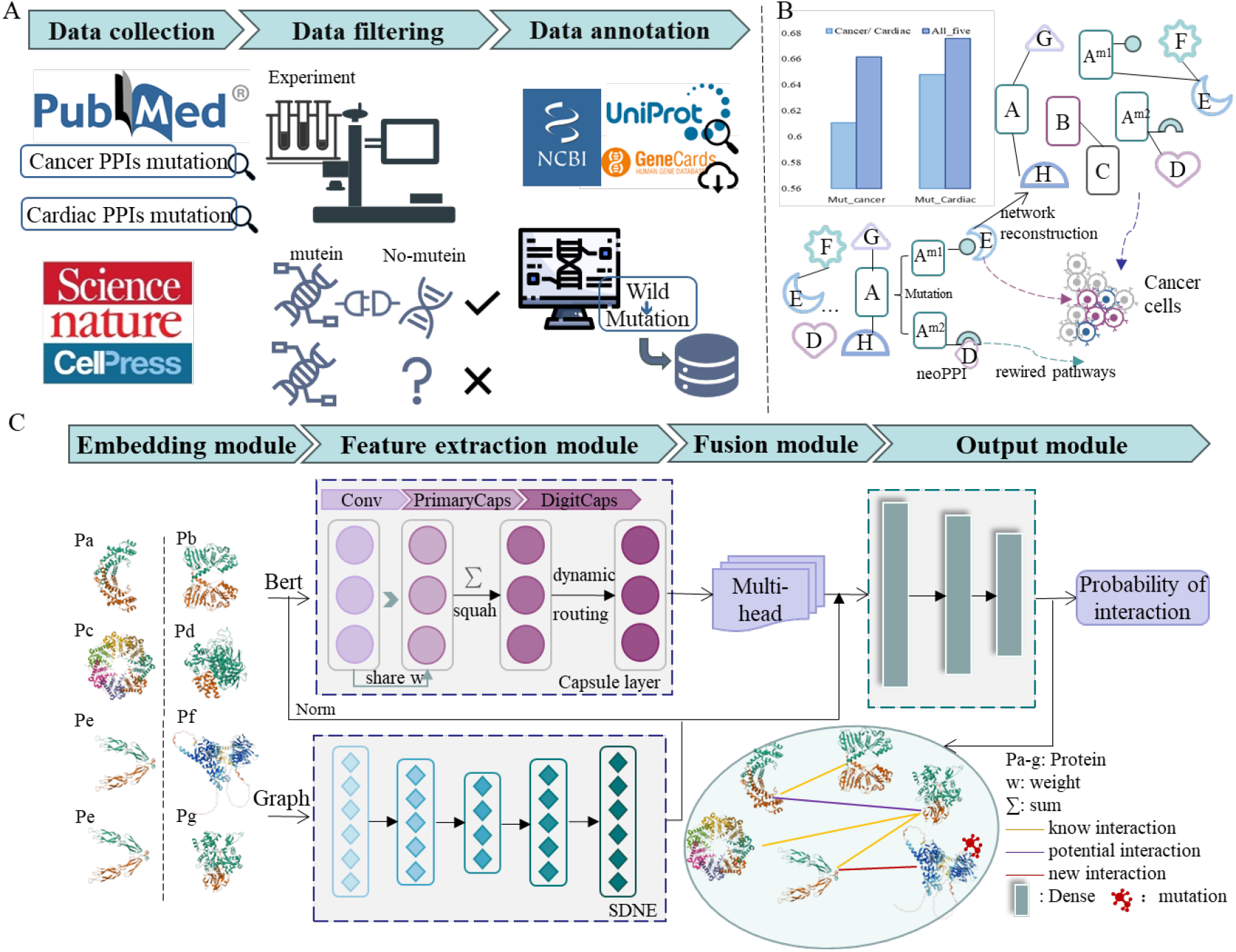
The overall work of the paper. A) Process for building a Real-world mutation-induced disease dataset using public database and resources. B) The mechanism of action of mutations leading to disease. C) The model of CGM architecture.

### 2.1 Real-world mutation-induced disease dataset construction

Currently, DL and AI methods are constrained by the quantity and quality of datasets. So, in this paper, constructing a high-quality real-world mutation-induced disease dataset is a critical step in predicting mutation-disease associations (Figure 1A). We focused on mutations in cancer and heart disease because more experimentally validated PPI data were available for these diseases. Querying PubMed (2021-2024) for experimentally determined PPIs (BRET, GST pulldown (PD), NanoPCA, semi-IP, affinity purification-mass spectrometry (AP-MS)) in cancer or heart disease returned 4 papers CITE, where one mutation was mutated and the interacting protein was not mutated.

We manually curated the data for validity, completeness, and experimental reproducibility. This yields a dataset of mutation, protein, and cancer or cardiac disease, encompassing 34 and 69 mutations, in 15 and 68 proteins in cancer and heart disease, respectively - Table 1 and see Supplementary Information. All data in this dataset have been obtained and validated through wet lab experiments such as affinity purification-mass spectrometry (AP-MS) ^30^ to ensure accuracy and reproducibility. Second, gene ID were converted to UniProt ID (using GeneCards ^31^, NCBI ^32^, and UniProt ^33^) and we converted the mutated proteins to their wild-type forms, and annotated them (Figure1 A).

**Table 1.** The performance and details of real-world mutation-induced disease dataset.

| Dataset | Interacting pairs | Proteins Number | Mutant Protein | Specific disease (prediction/all) | All_five (prediction/all) |
| --- | --- | --- | --- | --- | --- |
| Mut_Cancer | 511 | 334 | 34 | 312/511 | 338/511 |
| Mut_Cardiac | 71 | 115 | 69 | 46/71 | 48/71 |

Based on this process, we mined and curated data from papers in Science and other journals ^10, 30, 34, 35^ to construct the real-world mutation-induced disease dataset. It includes new protein interaction subsets related to cancer and heart disease, namely Mut_cancer and Mut_Cardiac (Table 1). These subsets contain 511 and 71 PPIs, involving 334 and 115 protein types, respectively. Detailed interpretations of the data are provided in the supplementary files.

### 2.2 Protein representation and Feature extraction

We designed the CGM to predict new PPI interactions where one protein is mutated (Figure 1B), which we can use to evaluate the impact of new protein interactions caused by mutations on disease (Figure 1C). To accurately characterize the interaction relationships of mutated proteins and other proteins in the PPI network, we used PPIs network for protein representation in the embedding module and introduced Bidirectional Encoder Representation from Transformers (BERT) to help overcome feature representation bottlenecks for protein sequences and highlight the features of mutated residues. Specifically, the multimodal representation includes extracting high-dimensional representation vectors using a BERT-based pre-trained protein model (Figure 2A) and converting the PPIs network into an undirected graph to obtain network structural information (Figure 2B). The feature extraction module designed different methods for the characteristics of embedded vectors. For high-level representations embedded by BERT, a capsule network was designed to capture contextual features, enhancing the association of high-level features obtained from the sequence. For network representation, SDNE was used to capture the nonlinear network information of the graph structure, ultimately producing a more compact representation. Then, the multi-head attention mechanism was employed to achieve information interaction and feature refocusing. Finally, the output module, a fully connected neural network layer, produces the probability of PPIs, which can determine whether the input protein pairs are likely to interact. The CGM not only utilizes BERT to refine the representation of protein sequence information but also attempts to construct a protein network to enhance the exploration of potential PPIs information from global data, providing opportunities to uncover new PPI relationships with the mutated protein. The multi-faceted embedding module lays a solid foundation for effective protein representation in mutation-oriented PPIs exploration.

**Figure 2.**
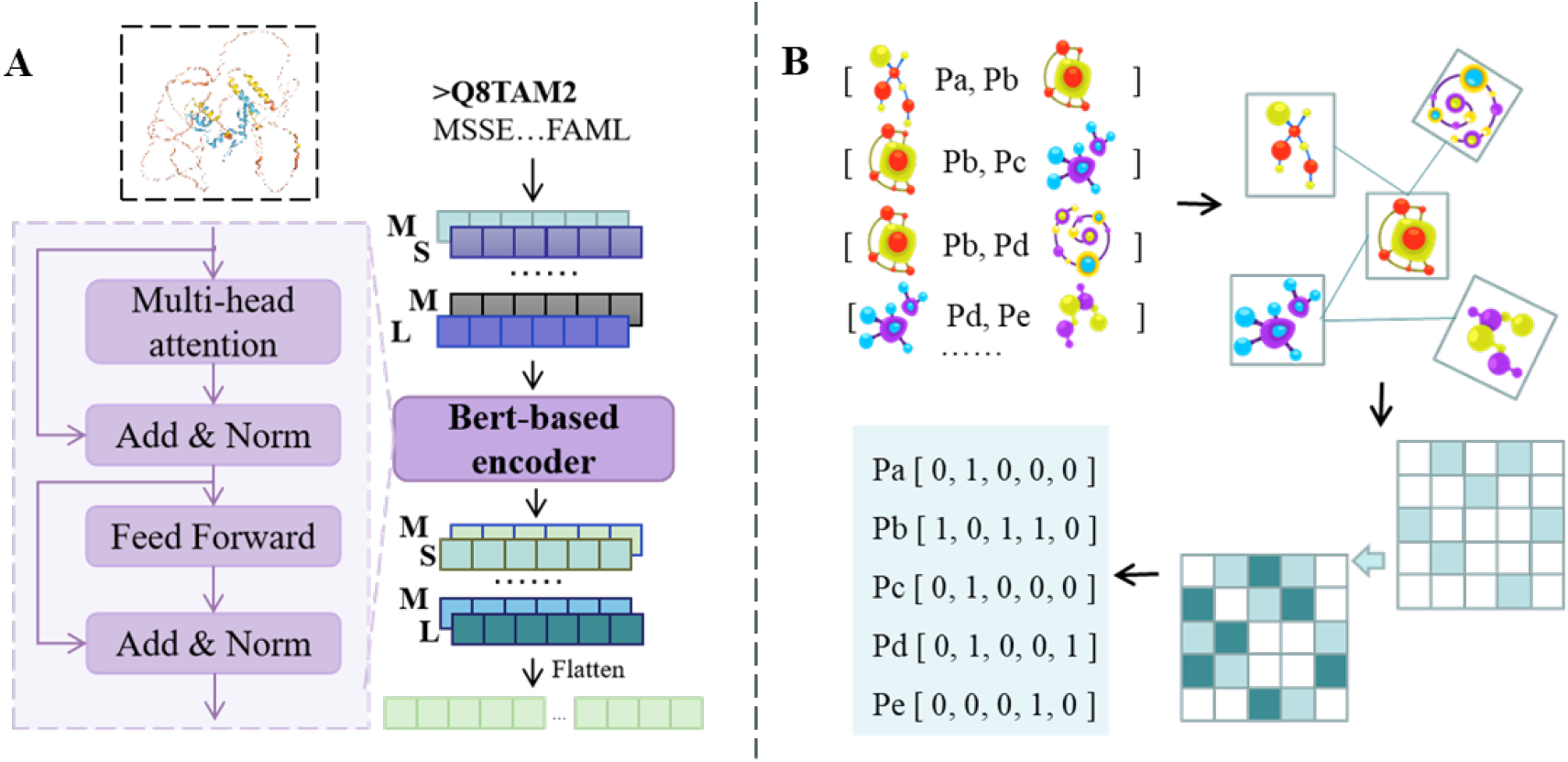
Details of the embedding module. A) Pre-training of the BERT model. B) The method allows to transform binary PPI into PPI graphs.

#### 2.2.1 Pre-training of the BERT model and capsule network

The pre-trained model is the BERT model based on a corpus of database of protein sequences^25^. This model can map protein residues into high-dimensional representations based on biological significance such as physicochemical properties, making it more sensitive to mutated residues. Specifically, this model is based on the general BERT framework and completes masked language modeling and next sentence prediction tasks on a corpus containing 2.1 billion protein sequences ^36^, ultimately achieving the mapping of input protein sequences to high-dimensional feature vectors. The details of the BERT model (Figure 2A) include multi-head attention mechanisms, fully connected networks, and residual connections. The multi-head attention mechanism, which integrates multiple independent self-attention modules, enables learning of multi-view contextual representations of protein sequences. The description of the multi-head attention mechanism is as follows.

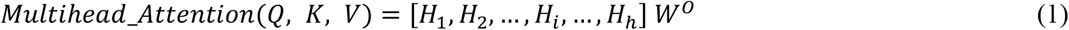

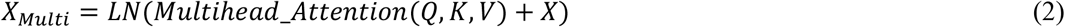

Here, *H*_*i*_ represents the *i-th* self-attention mechanism, and *W*_*0*_ establishes a mapping relationship between the output of the multi-head attention and the initial embedding dimensions. Then, residual connection techniques and layer normalization (LN) are used for deep feature extraction. The formula for *H*_*i*_ is:

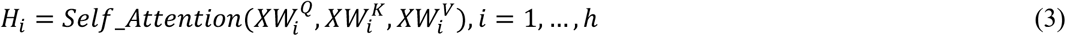

Where, 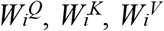 are the query, key, and value linear transformation layers of the *i-th* head, and *h* means the number of head.

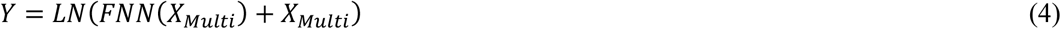

Subsequently, the synergy of residual connection techniques and layer normalization (LN) accelerates model convergence and improves overall performance. Finally, a fully connected network is used to better extract features. The mathematical formulas for this part are as follows.

Based on the high-dimensional protein sequence features encoded by BERT, which contain rich contextual semantic information, Capsule Networks ^37, 38^ can efficiently capture hierarchical features on a global scale due to their ability to analyze complex spatial hierarchical relationships. The principle behind this is that Capsule Networks replace individual neurons with vectors composed of groups of neurons, which together form capsules. Each layer of neurons in a Capsule Network contains multiple basic capsule units, which helps expand the receptive field of protein feature vectors. The neurons in Capsule Networks are connected in a manner similar to fully connected networks (Equations 5-10). This network includes three parts: Convolution Layer (Equation 5), Primary Capsules Layer (Equations 7-8), and Digit Capsules Layer (Equations 7-10). The Convolution Layer extracts local pattern features from high-dimensional representations, preparing for efficient hierarchical feature extraction. The Primary Capsules Layer aggregates local features from the Convolution Layer, while the Digit Capsules Layer performs overall modeling of the entire protein sequence using dynamic routing coefficients, with a calculation process similar to the previous layer.

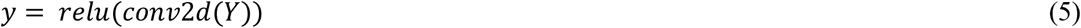

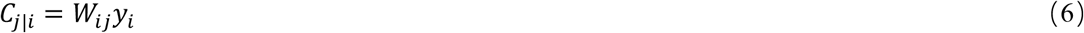

Where, *W*_*i*j_ means the weight matrix of the layer, and the *c*_*i*_ can describe the *i-th* layer of capsules. Then, each transformed vector is multiplied by a coefficient b_ij_ and passed to the next layer of capsules. Sum up all neuron signals received by the *j-th* capsule in the next layer (squah).

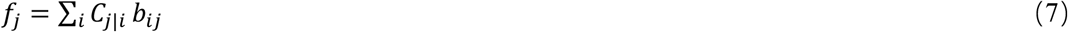

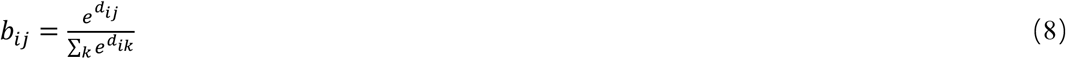

Here, *d*_*ij*_ is the logarithmic prior probability of whether two capsules are connected, and *k* represents the number of capsules in this layer. The output *pro*_*j*_ of this layer’s capsules can be represented by Equation 9, where *b*_*ij*_ starts with an initial value of 0 and is updated subsequently by a dynamic routing strategy (Equations 10).

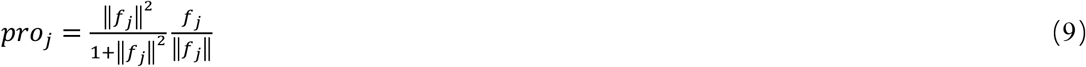

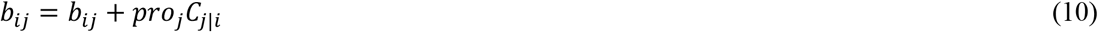

#### 2.2.2 The PPIs network and SDNE

When constructing a global relationship map of proteins based on the PPIs network ^27^, the PPIs network can be viewed as an undirected graph *G (N, E)*, where *N* represents the nodes of the graph, denoting proteins in this study, and *E* represents the edges of the graph, denoting relationships between proteins, that is, interaction or non-interaction. By including all proteins within the scope of mutation nodes, we can observe the connections between mutated proteins and potential proteins from a global perspective. Therefore,

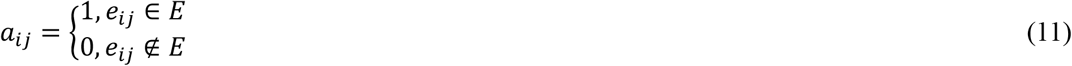

Among them, when *a*_*ij*_ is 1, it means that the two proteins connected by *e*_*ij*_ have interaction, and *a*_*i*_ is the adjacency matrix of *n*_*i*_, which describes the relationship between this protein and other proteins.

Structural Deep Network Embedding (SDNE) ^27^ could effectively capture highly nonlinear network structures. Therefore, we utilize SDNE to extract the nonlinear network features from PPIs graphs, generating denser representations. To preserve both local and global network structures, SDNE, like LINE (Large-scale Information Network Embedding), considers both first-order and second-order proximities. However, SDNE trains the first-order and second-order models separately and then embeds the results into the network, ultimately achieving a model that simultaneously considers both proximities for joint optimization. This joint optimization is accomplished through the mapping and folding of linear layers. By mapping and then remapping the original vectors to their restored dimensions, a more reasonable representation is achieved. The formulas are as follows:

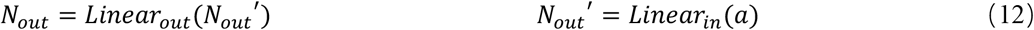

When *a*_*ij*_ is greater than 0, there is a positive first-order proximity between *n*_*i*_ and *n*_*j*_, mainly reflecting the local features of the graph. The second-order proximity indicates the similarity between two nodes, considering not only direct neighbors but also nodes with common neighbors. This means that structurally similar nodes can learn similar embedding vectors, hence allowing to capture global features through optimization. With the calculation of first-order and second-order similarities, the protein interaction information extracted from the network structure is more reasonable. To ensure structural authenticity, the corresponding local and global loss functions are as follows, where ⊙ denotes element-wise multiplication, and *c* is a hyperparameter.

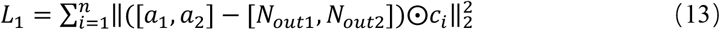

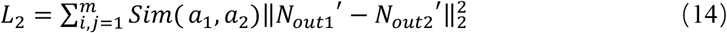

Where *a*_*n*_ is the adjacency matrix vector of node *n*, and the structural similarity value is calculated by the following formula:

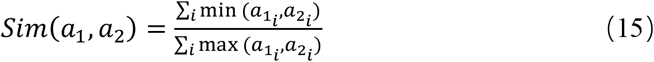

### 2.3 Fusion and Output module

Typically, PPIs calculation methods concatenate the protein representations rigidly before proceeding to downstream tasks, which can lead to information redundancy and diminish the possibility of generate new information. Therefore, based on the biological similarity between protein pairs, multiplying the query information with other proteins can enhance the interaction between the predicted proteins, strengthening the exploration of crucial information between protein pairs.

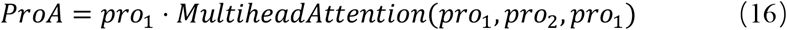

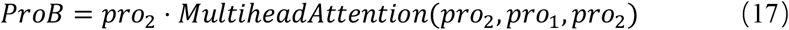

Here, *pro*_*1*_ and *pro*_*2*_ are the feature vectors of the protein passing through the capsule network. The final protein expression is then obtained by global max pooling as follows:

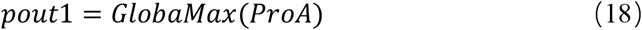

Correspondingly, *ProB* also produces an output, which is *pout2*. The skip connections allow the input of a layer to be bypassed through one or more layers and directly added to the output of subsequent layers. This helps preserve important information, enhance feature propagation, and better capture long-range dependencies, thereby promoting the training of deeper networks. Therefore, by employing skip connections, we enhance the data processing ability of the network and reduce information loss during feature extraction. Additionally, to strengthen the positional correlation and interpretability between protein information pairs, we incorporated cosine similarity (cos_sim_), bilinear similarity (BiLinear_sim_), and linear similarity calculations (Linear_sim_), concatenating these attribute features as follows:

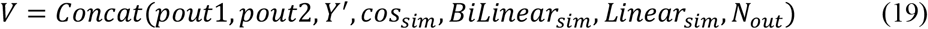

Where *Y’* is the vector normalizing *Y*. The output module is to perform multiple linear transformations on the fusion module (Equation 19), and finally obtain the final output result through the sigmoid function. Where *w*_*i*_ is the contribution weight of the output and w_0_ is the additive bias term.

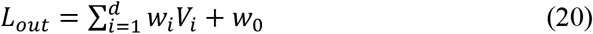

Simultaneously, the model utilizes two feature extraction modules, capsule networks and SDNE, to handle the obtained high-dimensional and network feature. In order to preserve the integrity of the network structure and the accuracy of the prediction task, we redefine the loss function of the model. However, the loss function of the capsule network is not accounted for because the reconstruction loss used has a negligible impact on performance ^37^.

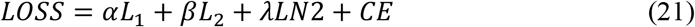

Where α, λ, and β are hyperparameters, LN2 is the L2-norm regularize added to prevent overfitting, and CE is the cross-entropy loss between the predicted and true values.

## 3 RESULTS

To achieve efficient mutation-disease association prediction, in this paper we first construct a real-world mutation-induced disease dataset from published, experimentally determined and manually curated mutated-proteins interactions, which is then used in prediction model for mutation-directed protein interaction. Specifically, we developed a mutation-disease associations prediction model based on CGM to assess the impact of mutations on novel protein interactions and disease. We also explore if altered protein structure due to mutations can cause disease and find that mutations can alter the protein structure and that different mutation sites on the same protein can lead to different structural changes. Although we constructed the real-world mutation-induced disease dataset, it is limited to the scale of mutated protein data and does not fully showcase the potential of CGM in predicting mutation-disease associations.

Given the guidance provided by discovering new protein interactions in existing disease data for mutation-guided new protein interactions, we also conducted experiments on mining unknown protein interaction pairs. And our results showed that CGM identified 22 unknown protein interaction pairs across benchmark datasets for diabetes, cardiac disease, Alzheimer’s disease (AD), Parkinson’s disease (PD), and cancer. We also compared CGM with other advanced methods using imbalanced datasets with varying positive-to-negative sample ratios. In most cases, CGM exhibited optimal results, further demonstrating its potential in discovering novel protein interactions related to function and disease. Detailed evaluation metrics for all experiments are provided in the supplementary files.

### 3.1 Mutation directed protein interactions performance

The models for PPIs prediction have traditionally focused on exploring protein relationships without attempting to explain the mechanisms of diseases caused by systemic disruptions due to protein mutations. The lack of data on mutation-directed protein interaction and limitations in model performance are the main reasons. Therefore, we designed the CGM model to discover mutation-induced PPIs in cancer and Cardiac. As shown in Table 1, the model correctly predicted mutation-induced novel PPIs in 312 out of 511 cases (61%) in the cancer-specific dataset and 46 out of 71 cases (65%) in the cardiac-specific dataset. Additionally, to test if the predicted mutated-protein interactions reflect similar disease mechanisms, we conducted prediction tasks across all disease datasets (all_five). The results indicated that more PPIs could be predicted in the all_five datasets, suggesting a degree of similarity in disease mechanisms. This viewpoint is reflected in the data from different diseases, for instance, the protein interaction pair Q13485 (SMAD4) and Q15796 (SMAD2) in the cancer dataset also appeared in the Cardiac dataset. Notably, Q13485 (SMAD4) and Q15796 (SMAD2) proteins are associated with Cardiac and tumors (https://www.uniprot.org).

We also analyzed the similarities between different disease datasets (Figure 3), further illustrating the prediction of common mutated-protein interactions across of diseases^39^. The most notable overlap is between PD and AD, sharing a significant number of PPIs (over 1/8 of positive samples), which is not surprising given that known relationship between these neurodegenerative diseases (PMID: 33154531). The secondary significant overlap is between PD and cancer, which is in keeping with a suggested link between PD and cancer^40^.

**Figure 3.**
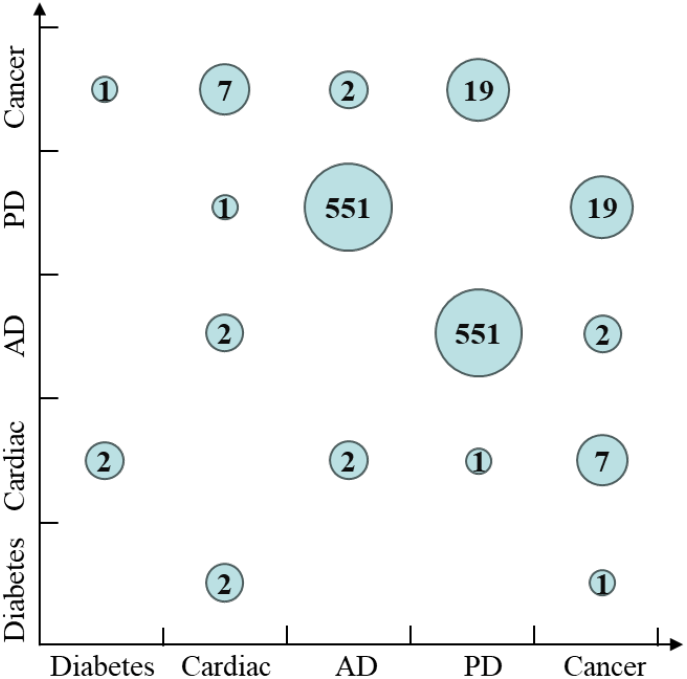
Statistics of the number of similarities in different disease datasets

To further explore the causal associations between mutated proteins and diseases, we examined the structural differences between mutated proteins and their wild types. Specifically, using the Swiss-model ^41^, we simulated the structures of wild protein and mutated proteins (Figure 4). For example, observing the mutated proteins P42336_E545K (PIK3CA_E545K) and P42336_K111E (PIK3CA_K111E), we found that the three-dimensional conformations of wild-type and mutated proteins differ, especially at the mutation sites. In Figure 4, the red box indicates the differences between wild-type proteins and mutant proteins. This mutation-induced structural changes might be a key factor in their interactions with other proteins, leading to disease.

**Figure 4.**
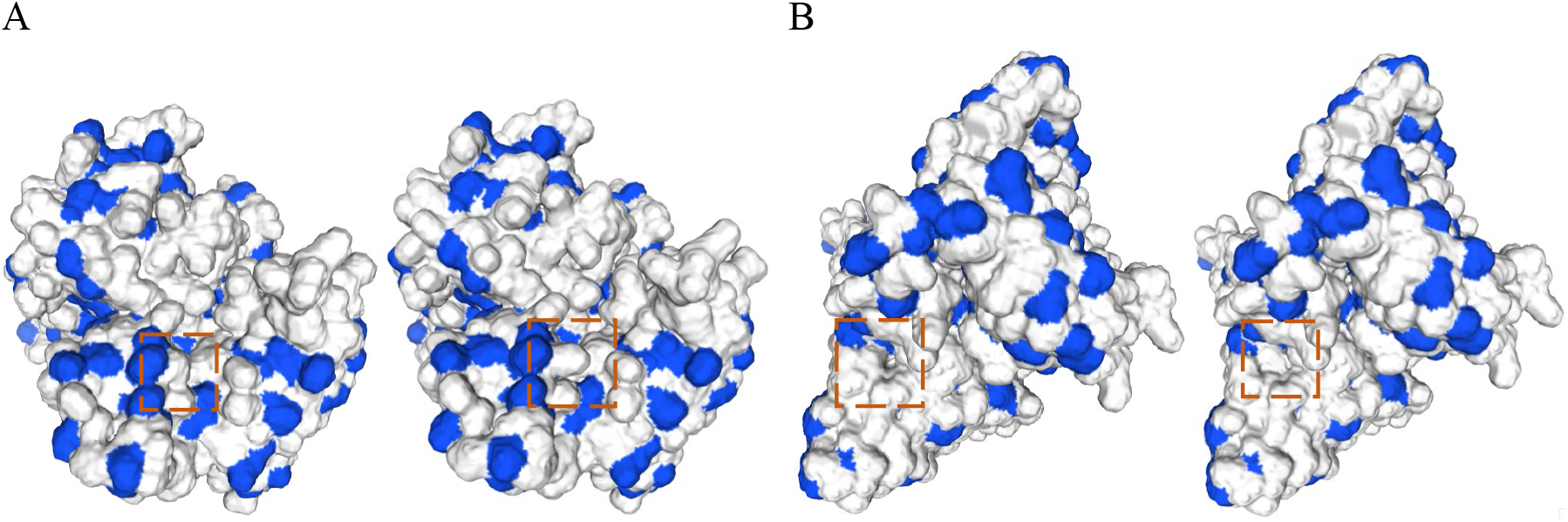
Comparison of 3D conformations between wild-type and mutated proteins. A) Conformation comparison between PIK3CA and PIK3CA_E545K. B) Conformation comparison between PIK3CA and PIK3CA_K111E

### 3.2 Analysis in Mining unknown PPIs and unbalanced datasets

Despite constructing the real-world mutation-induced disease dataset, limited by the data and wet-lab experiment scale constraints, it does not fully showcase the potential of CGM in predicting mutation-disease associations. Therefore, this section we primarily illustrate the ability of the CGM model to predict PPIs within disease datasets, and investigate previously unknown protein interactions, predicted by our model. Since in real case applications, there are typically fewer samples of disease-induced mutated PPIs (positive samples), with negative samples being more readily available ^42^. This imbalance in the dataset can severely impact model performance. Therefore, we analyzed the performance of CGM on imbalanced datasets.

Five publicly accessible datasets ^43^ were retrieved from DIP and IntAct databases (as shown in Table 2), covering Diabetes, Cardiac, AD, PD, and Cancer, and combined into a merged dataset (All_five). Protein sequence information is taken from the UniProt platform ^33^.

**Table 2.** Details of the interaction datasets.

| Dataset | Interacting pairs | Noninteracting pairs | Number of proteins |
| --- | --- | --- | --- |
| Diabetes | 163 | 163 | 121 |
| Cardiac | 2663 | 2663 | 1840 |
| Alzheimer disease (AD) | 4096 | 4096 | 2866 |
| Parkinson's disease (PD) | 4127 | 4127 | 3271 |
| Cancer | 6352 | 6352 | 3909 |
| All_five | 17401 | 17401 | 8478 |

#### 3.2.1 Exploring potentially unknown PPls

In terms of verifying the application potential of the model, we found several protein pairs with potential new interactions based on the all_five dataset (Figure 5 and Supplementary File S2). We used Cytoscape^44^ to create Figure 5, where the nodes represent proteins, and different colors indicate data from various disease datasets. Solid lines represent interactions from the original network, while dotted line represent previously unknown, and potentially novel, protein interactions in the dataset. Thicker edges indicate higher similarity. Figure 5 demonstrates that the model captures the potential PPIs not mentioned in the dataset. And these PPIs are input into STRING 45 to further provide the basis for the applicability of the model. Through literature mining, gene co-expression, and other methods, it is found that the unknown PPIs predicted by CGM are adaptive, which proves the rationality and interpretability of the model. For instance, P49207 (RPL34) ^46^ and O60885 (BRD4) ^47^ play crucial roles in cancer development and progression. The connection between them in STRING indicates a high probability of interaction, supported by experimental evidence ^48, 49^ and enhanced by co-expression and data mining. Other interacting protein pairs can also find correspondence descriptions in the STRING dataset to illustrate that CGM is able to discover PPIs that may have interaction relationships.

**Figure 5.**
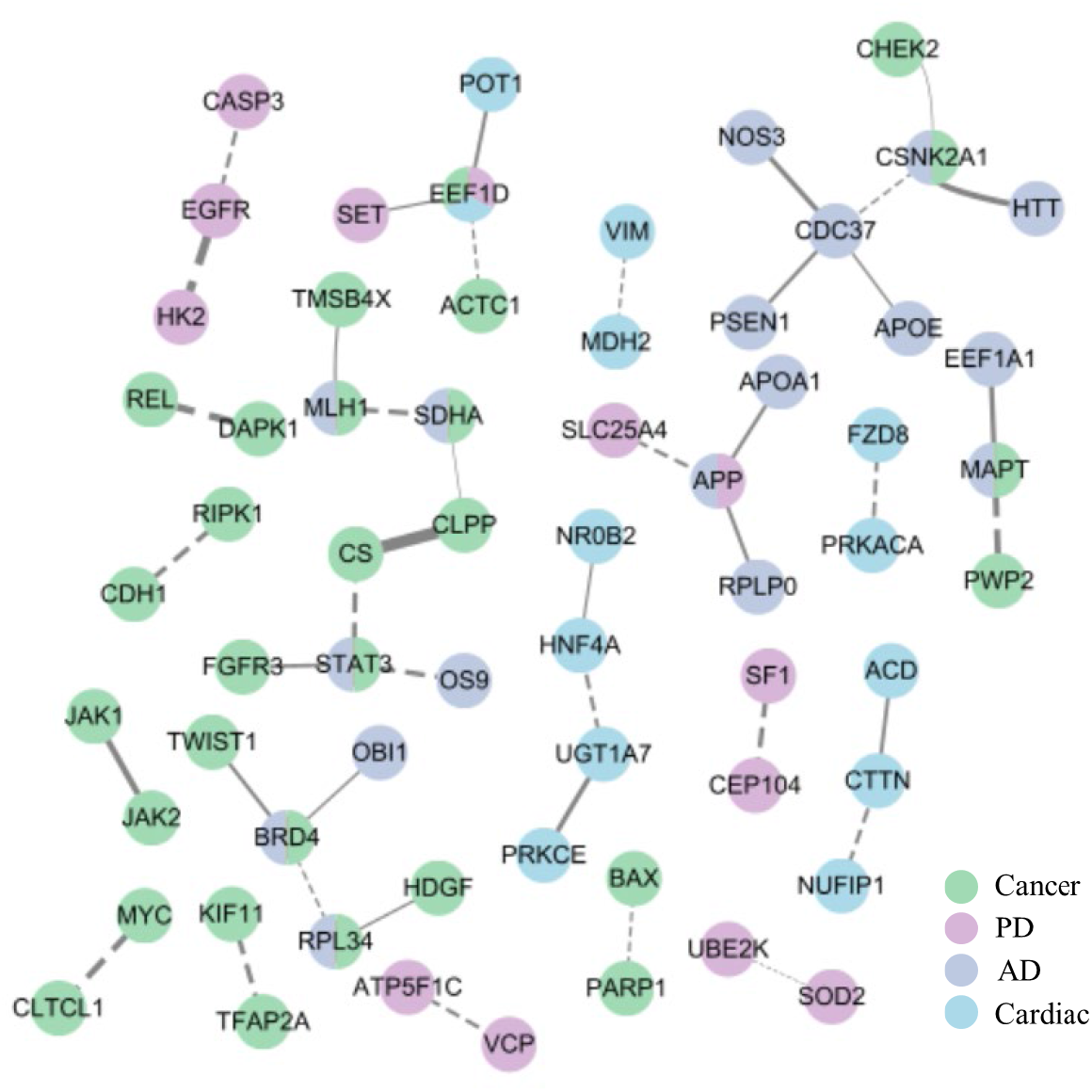
The unknows PPIs (dotted line) captured by the CGM that are not previously reported. Continuous line indicates known PPI.

#### 3.2.2 Performance of different proportions of imbalanced disease datasets

Considering that in practical applications, the number of positive disease samples is often less than negative samples, this section compares CGM with other advanced method on datasets with different sample ratios to further analyze the performance of CGM in mutation tasks. Table 2 shows balanced positive and negative sample data, resulting in low sensitivity of the model to the small number of positive samples. Therefore, we adjusted datasets with different ratios of PPIs and compared CGM with the SGAD method. The results indicate that CGM outperformed SGAD in most metrics (46/50) (Supplementary Tables S3-S4). Additionally, we conducted a comprehensive analysis of multiple imbalanced datasets, as shown in Figure 6, which uses error bars to display the observed variations in the imbalanced datasets.

**Figure 6.**
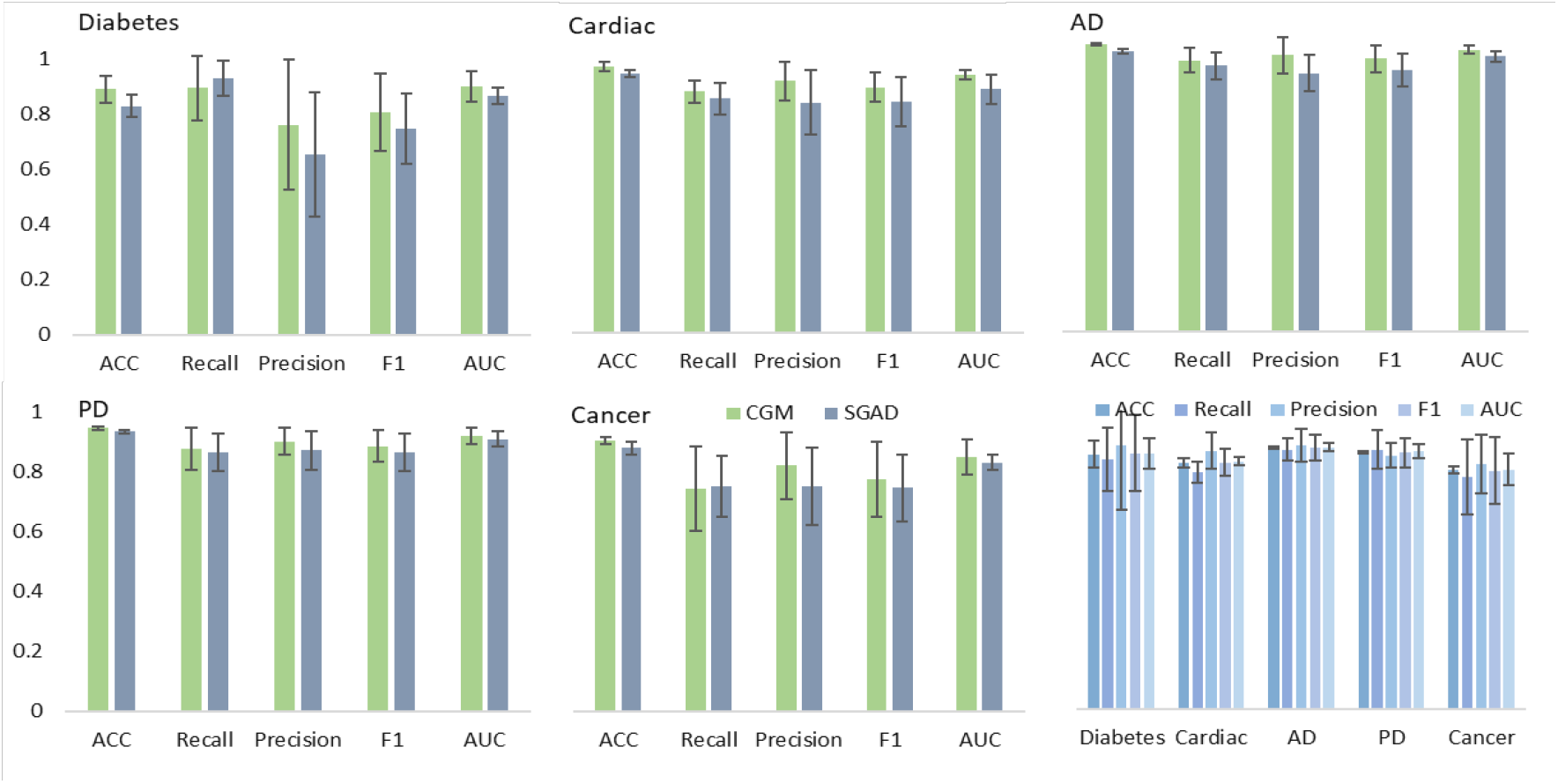
Presentation of the mean and error bars for different datasets at different radios

The Figure 6 shows that the average values of CGM in three imbalanced datasets are generally higher than those of SGAD (24/25), and more than half of the error bars are shorter than those of SGAD. This demonstrates that CGM has less fluctuation in evaluation values on imbalanced datasets, showing higher stability in predicting disease-related PPIs caused by mutations, even with few positive samples. We also analyzed performance across five disease datasets and found that the average performance on the small sample dataset (diabetes) was comparable to other datasets, although stability still needs improvement.

### 3.3 Comprehensive Performance Comparison

To further demonstrate the potential of CGM in PPIs, we compared CGM with other advanced baseline models on benchmark datasets. The results indicate that CGM not only identifies unknown protein interactions in disease datasets and exhibits excellent performance on imbalanced datasets, but also achieves good results in benchmark tasks. Comparison tasks are designed based on five datasets (Diabetes, Cardiac, AD, PD, and cancer) to show the prediction performance of CGM and other reported methods, including NDDs_APAAC ^50^, EnsAmDNN ^50^, EnsDNN ^50^, SGAD ^27^, and MARPPI ^26^. The evaluation metrics consistent with the compared methods include accuracy (ACC), recall, precision, F1-score, and Area Under the ROC Curve (AUC), with values closer to 1 indicating superior performance (detailed interpretations of the evaluation metrics are provided in the supplementary files).

Figure 7 shows the performance results of the models on different disease datasets. The results indicate that CGM shows significant advantages in various evaluation metrics across the five datasets (Supplementary Files S5-S10). In the diabetes, the CGM achieved an accuracy close to 0.94, improving by 0.05-0.14 over other methods, and F1 improved by 0.04-0.15. Additionally, compared to the MARPPI method, which employs pre-trained models and descriptor feature embedding, CGM outperforms in metrics like accuracy, highlighting the advantages of our feature embedding approach. We also observed significant improvements of at least 0.03 in multiple metrics for cancer and AD. Although CGM did not achieve the highest values in all metrics for the cardiac, it was closest to the highest values, with a difference of only 0.016. From a comprehensive evaluation perspective, the method still shows an advantage, though further improvements in precision are needed. Summarizing the five disease datasets, we found that the cancer dataset metrics were generally lower than those of other datasets, possibly due to the higher diversity of proteins (Table 2), making the network characteristics more complex and challenging to capture.

**Figure 7.**
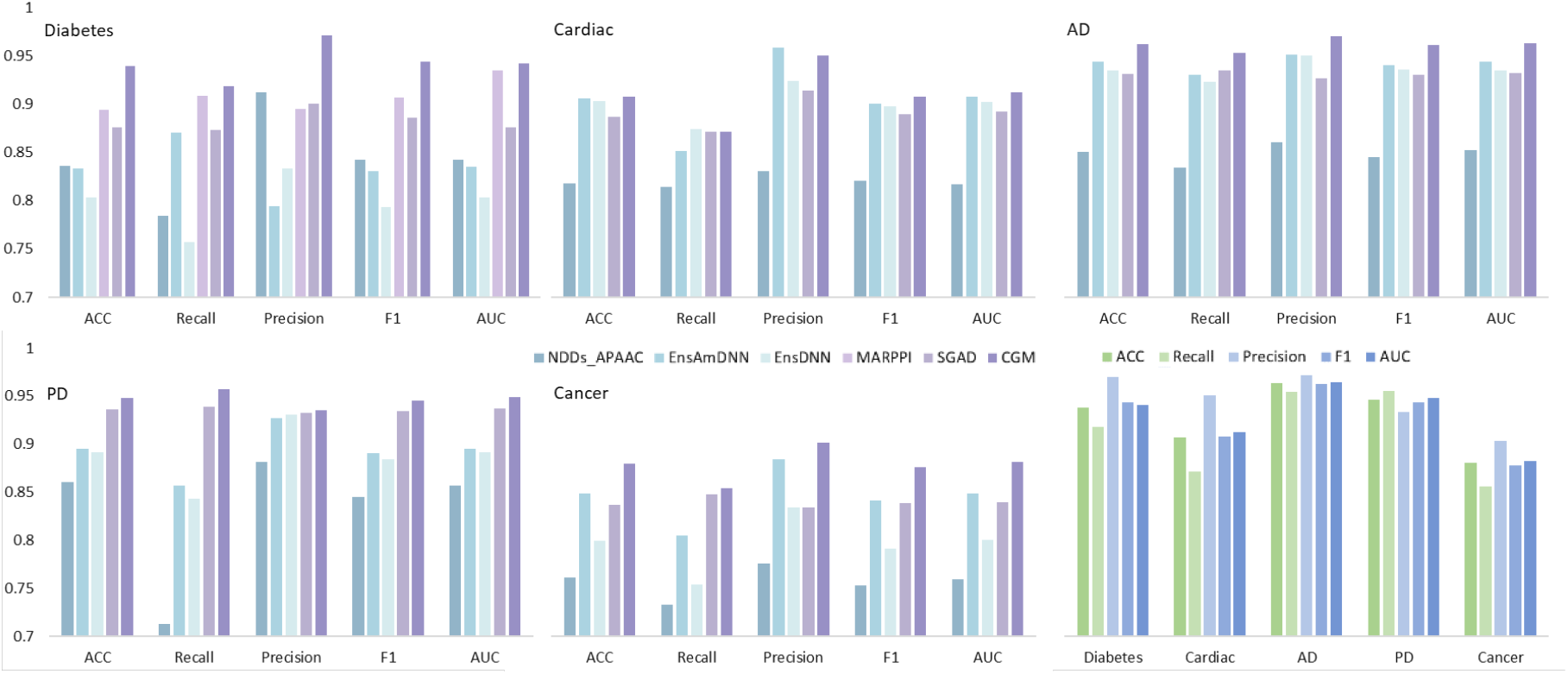
Comparison results of different prediction methods on the different datasets

### 3.4 Ablation experiments

To assess the effectiveness of each component in the CGM model, we conducted a series of ablation experiments based on the diabetes dataset (Table 3). We designed a model (model_0) that replaces the capsule network with a linear layer, which also lacks the PPIs network and the information interaction module. According to the description, CGM can alleviate the rigid concatenation of two protein vectors and facilitate information interaction. Therefore, model_1 is a model without protein information interaction and also lacks the PPI network. The results of model_0 and model_1 demonstrate the advantage of capsule networks in feature extraction. Based on the advantages of the CGM model in predicting unknown protein-protein interactions (PPIs) induced by mutated proteins, we designed model_2, a model without the PPI network. The results of model_1 and model_2 indicate the necessity of information interaction, which can reduce information redundancy while improving accuracy. Compared with model_2, CGM includes the PPI network module. The results show that the presence of the PPI network can extract the structural features of the protein network globally. Although it increases the feature dimensions, it enhances learning depth and improves model performance.

**Table 3.**
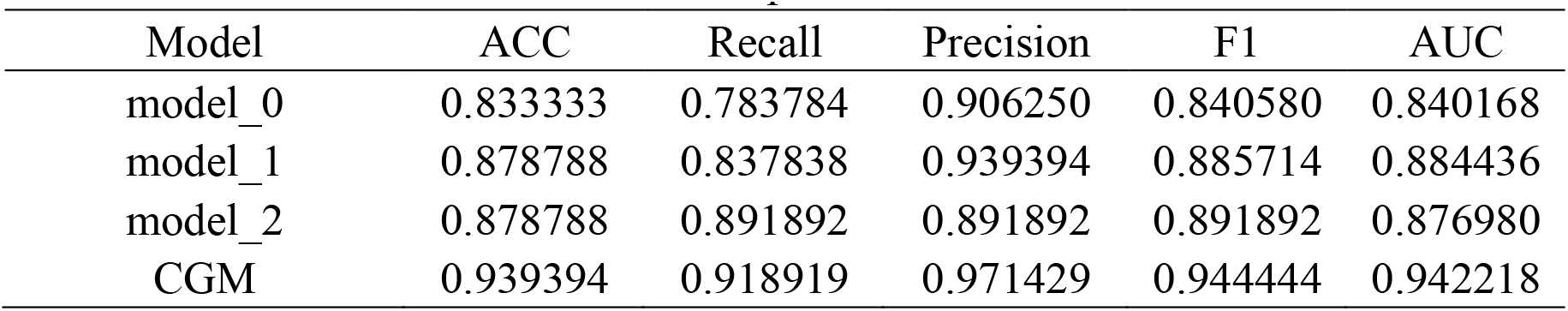
Performances of the different component.

| Model | ACC | Recall | Precision | F1 | AUC |
| --- | --- | --- | --- | --- | --- |
| model_0 | 0.833333 | 0.783784 | 0.906250 | 0.840580 | 0.840168 |
| model_1 | 0.878788 | 0.837838 | 0.939394 | 0.885714 | 0.884436 |
| model_2 | 0.878788 | 0.891892 | 0.891892 | 0.891892 | 0.876980 |
| CGM | 0.939394 | 0.918919 | 0.971429 | 0.944444 | 0.942218 |

## 4 CONCLUSIONS

This paper constructs a real-world mutation-induced disease dataset and designs a prediction method called CGM to predict on mutation-based protein interactions. To accurately characterize the interaction relationships of mutated proteins within the protein network, we use the PPI network structure in the embedding module and introduce BERT to capture high-potential representations of residue contextual information in the sequence, enhanced feature representation of protein. At the same time, we use the capsule network and multi-head attention mechanism to capture high-latent information to complete information interaction, strengthen local information, and finally form a dependency relationship with the protein network from both local and global perspectives to efficiently complete the prediction task.

The results of the constructed and benchmark datasets illustrate that CGM not only achieved desirable results on the mutation-disease association prediction task but also mined 22 previously unknown protein interaction pairs. In particular, datasets with different sample proportions, the CGM shows over 84% accuracy and consistency at different datasets, validating the robustness and stability. To further investigate the pathogenic mechanisms of mutated proteins, we explored structural changes based on the theory that protein mutations affect structure, which in turn affects disease. The results demonstrate that protein mutations can cause structural changes, and different mutation sites lead to different structural changes. And the results were demonstrated to be consistent with biological phenomena, highlighting a certain level of biological analysis capability that can be used to explore unknown protein pairs and mutation-based protein interactions related to diseases.

In summary, this paper uses deep learning to predict the protein mutations-disease associations, providing a novelty tool for further understanding biological molecular pathways and disease mechanisms. Although the CGM model shows excellent performance in predict mutations-disease associations, the current mainstream view primarily considers diseases to be strongly related to changes in protein function^51^. Therefore, our next primary task is to further analyze functional changes in proteins post-mutation. Additionally, the scale of the mutation-causing disease dataset needs to be further expanded, serving as foundational work for deeply analyzing the pathogenic mechanisms of proteins.

## ASSOCIATED CONTENT

### Data Availability Statement

The data utilized in this study were sourced from publicly available datasets, accessible through the following websites: https://www.ebi.ac.uk/intact, https://www.genecards.org/, https://www.uniprot.org/, and https://string-db.org/. The source code for the implementation of the methods discussed in the article is available at https://github.com/xueleecs/CGM.git.

## Author Contributions

X.L. and B.C. collected data, developed the model, analyzed the data; J.W., X.M., S.W., and Y.H. helped to refine the research through constructive discussions and revised the manuscript; E.P. and T.S. supported and supervised the project, interpreted the results, and wrote revisions to the manuscript.

## Funding Sources

This work was supported by the National Natural Science Foundation of China (Grant Nos. 62272479, 62202498, 62372469), Taishan Scholarship (tsqn201812029), Foundation of Science and Technology Development of Jinan (201907116), Shandong Provincial Natural Science Foundation (ZR2021QF023), Spanish project (PID2019-106960GB-I00), and Juan de la Cierva (IJC2018-038539-I), China Scholarship Council.

## Notes

The authors declare no competing financial interest.

## Acknowledgements

We would thank Jing Guo and Yinuo Zhang for their insightful discussions.

## Notes

### Competing Interest Statement

The authors have declared no competing interest.

